# Small protein mediates inhibition of ammonium transport in Methanosarcina mazei – an ancient mechanism?

**DOI:** 10.1101/2023.09.04.555848

**Authors:** Tim Habenicht, Katrin Weidenbach, Adrian Velazquez-Campoy, Ruben M. Buey, Monica Balsera, Ruth A. Schmitz

## Abstract

In the past decade, small open reading frames (sORFs) coding for proteins less than 70 amino acids (aa) in length have moved into the focus of Science. sORFs and corresponding small proteins have been recently identified in all three domains of life. However, the majority of small proteins remain functionally uncharacterized. While several bacterial small proteins have already been described, the number of identified and functionally characterized small proteins in archaea is still limited. In this study, we have discovered that the small protein 36 (sP36), which consists of only 61 aa, plays a critical role in regulating nitrogen metabolism in *Methanosarcina mazei.* The absence of sP36 significantly delays the growth of *M. mazei* when transitioning from nitrogen limitation to nitrogen sufficiency, as compared to the wild type. Through our *in vivo* experiments, we have observed that during nitrogen limitation, sP36 is dispersed throughout the cytoplasm; however, upon shifting the cells to nitrogen sufficiency, it relocates to the cytoplasmic membrane. Moreover, in vitro biochemical analysis clearly showed that sP36 interacts with high-affinity with the ammonium transporter AmtB_1_ present in the cytoplasmic membrane during nitrogen limitation, as well as with the PII-like protein GlnK_1_. Based on our findings, we propose that in response to an ammonium up-shift, sP36 targets the ammonium transporter AmtB_1_ and inhibits its activity by mediating the interaction with GlnK_1_.

**Importance:** Small proteins containing fewer than 70 aa, which were previously disregarded due to computational prediction and biochemical detection challenges, have gained increased attention in the scientific community in recent years. However, the number of functionally characterized small proteins, especially in archaea, is still limited. Here, by using biochemical and genetic approaches, we demonstrate a crucial role for the small protein sP36 in the nitrogen metabolism of *M. mazei*, regulating the ammonium transporter AmtB_1_ according to nitrogen availability. This regulation might represent an ancient archaeal mechanism of AmtB_1_ inhibition by GlnK, in contrast to the well-studied regulation in bacteria, which depends on covalent modification of GlnK.

## Introduction

Small proteins are a significant part of the proteome in all organisms across the tree of life. The definition of this class of proteins is based on their small size rather than a functional criterion, which is typically defined by a cut-off ranging from 70 to 100 amino acids ^1–5^. Unlike peptides produced through post-translational processing of larger proteins, the small proteins we refer to here are encoded by independent small open reading frames (sORFs). These sORF-encoded proteins, with a length of fewer than 70 amino acids (aa), have been historically overlooked and understudied due to bioinformatic biases inherent in conventional genome annotations, and technical challenges associated with classical biochemical approaches, such as SDS-PAGE or mass spectrometry. Conventional gene annotations were primarily designed to identify larger proteins^6,7^, while proteomic tools relied on obtaining multiple peptides of one protein through tryptic digestion, which is often not feasible for small proteins^8^. These challenges have impeded the annotation and characterization of small proteins in the past, creating a promising avenue for detailed mechanistic studies and functional analysis today, with high untapped potential.

Genome-wide transcriptomic, translatomic and proteomic methods have been improved and developed in recent years to address and overcome the challenges to identify the small proteome. The application of deep sequencing technologies, as well as improvements and adaptation of ribosome profiling tools to bacteria and archaea, and optimized peptidome analyses by mass spectrometry allowed the identification of a constantly growing number of sORFs and the respective small proteins in bacteria and archaea^9–14^. As a result, an increasing number of reports on small proteins encoded by sORFs are currently emerging and their physiological importance has been proven in numerous examples, by participation in various cellular functions such as cell division, transport and enzymatic processes^3,15,16^.

Since small proteins have come in the focus of Science, it becomes more evident that a significant amount of proteins with less than 70 aa is associated with the cytoplasmic membrane. For *E. coli*, a large portion of identified small proteins were shown to be localized at the membrane, where they might interact with larger proteins and protein complexes such as signal receptors or transporters^17–19^. However, archaea exhibit fewer identified small proteins, and functional characterization is limited to a handful of examples, of which only one is part of a transporter^20–23^.

*Methanosarcina mazei* strain Gö1 belongs to the order Methanosarcinales and is strictly anaerobic. This versatile group of methylotrophic archaea can utilize a variety of substrates, including methanol, methylamines, acetate, in addition to CO_2_ and H_2_, as a source of both carbon and energy, with the ultimate product being methane, a greenhouse gas^24,25^. In the absence of another suitable nitrogen source, *M. mazei* is able to reduce and fix molecular nitrogen^26^. This highly energy consuming process of nitrogen fixation as well as the general nitrogen metabolism are strictly regulated on transcriptional, posttranscriptional and posttranslational level in response to nitrogen availability, which has been studied extensively in recent years^27–33^. As known for bacteria, several key components of the nitrogen metabolism are only present and highly expressed under N-starvation e.g. glutamine synthetase, ammonium transporters and in diazotrophs also nitrogenase^32,34–37^.

Under sufficient ammonium concentration, nitrogen assimilation in cells occurs through the diffusion of ammonia across the cytoplasmic membrane and its subsequent incorporation into glutamate by glutamate dehydrogenase. However, under conditions of significantly decreased external ammonium concentration, active transport of ammonium becomes necessary. This transport is facilitated by the trimeric ammonium transporter AmtB_1_, which is expressed exclusively under nitrogen (N) limitation in *M. mazei*^10,38^ but requires the expenditure of ATP^39,40^. Following transport, ammonium assimilation is facilitated by the glutamine synthetase/GOGAT system^41^. N starvation leads to elevated levels of 2-oxoglutarate (2-OG) within cells, which serves as an internal signal for N starvation^32^.

AmtB_1_ represents an ammonium transporter protein of the Amt/Mep/Rh protein family, of which, members can be found in eukaryotes, bacteria as well as in archaea. Through all domains of life, proteins of the Amt family show a highly conserved tertiary structure of 11 transmembrane helices with extracellular N- and cytoplasmic C-terminal domains^42,43^. In *Escherichia coli*, the ammonium transporter AmtB organizes as trimers with each subunit representing a hydrophobic pore for ammonia transport^44–46^. The import of NH_4_ ^+^ is an energy consuming process^39,40^. Consequently, the transporter is highly regulated depending on the nitrogen status of the cell to exclude energy dissipation. Based on structure and complex formation analysis, Coutts et al.^46^ showed that in *E. coli* the ammonium transporter is inhibited by a PII-like protein (GlnK) upon a shift to nitrogen sufficiency. This regulation is based on a GlnD-dependent deuridylylation of GlnK. The respective *glnK* gene is organized together with the *amtB* gene in an operon (glnK/amtB), which is only expressed under nitrogen starvation^47^. This coupling of genes encoding an ammonium transporter and a PII-like protein has been identified in most bacteria as well as in archaea indicating a tight functional coexistence^48^.

A large number of potential sORFs were identified in *M. mazei* under N stress conditions through genome-wide RNAseq analysis^10^. Among these, sORF36 encodes a 61 aa protein (sP36) that was confirmed by LC-MS/MS analysis. Its transcription was shown to increase 2.5-fold under nitrogen limitation, as confirmed at the protein level. Additionally, sORF36 and sP36 are highly conserved on the DNA and protein level, across various Methanosarcina strains, suggesting a possible role of sP36 in nitrogen metabolism^31^.

In this study, we characterize this additional newly discovered component in nitrogen regulation in *M. mazei*, the small protein sP36^31^. Through genetic and biochemical approaches, we show that sP36 is involved in the adaption to changing nitrogen conditions. Although sP36 does not contain any transmembrane helices, we provide several lines of evidence *in vitro* and *in vivo*, that sP36 localizes at the cytoplasmic membrane in response to an ammonium upshift after a period of nitrogen limitation. Using different biochemical approaches, we demonstrated that the observed interaction with the cytoplasmic membrane is the result of the direct interaction between sP36 and the membrane located ammonium transporter AmtB_1_. In a pull-down assay, purified His_6_-tagged AmtB_1_, incubated with native *M. mazei* cell extract, is capable to specifically mediate the retention of chromosomally expressed sP36. Further biochemical analysis demonstrated high affinity interaction between sP36 and not only the ammonium transporter AmtB_1_ but also with the regulatory PII like protein GlnK_1_.

We propose a plausible model where sP36 acts as an adaptor protein that mediates the interaction GlnK_1_-AmtB_1_ to allow a rapid and reversible response to changes in nitrogen availability. This mechanism might represent a more ancient version of the AmtB inhibition by a PII-like protein before the GlnD-dependent uridylylation was developed.

## Results

### sP36 plays a crucial role during nitrogen upshifts after a period of N limitation

To get insights into the functional role of sP36 a genetic approach was performed. A chromosomal deletion mutant of the respective sORF encoding sP36 was constructed, replacing the *sORF36* gene with the puromycin resistance cassette (*pac* cassette) using an allelic replacement approach (see MM). The generated mutant strain (*M. mazei* ΔsP36) was verified by Southern blot analysis (Fig. A1). Its growth behavior under different N availabilities was evaluated and compared to the wild type strain (*M. mazei* 3A; Fig. 1). When growing on minimal medium with ammonium as sufficient N source (10 mM) or under N limitation (0 mM) no growth phenotype was detectable in the absence of sP36 except that the lag phase was slightly prolonged but reaching identical doubling times. However, when cells are grown under N limitation until early exponential phase (turbidity at 600 nm (T_600_) = 0.15) and then transferred into fresh ammonium sufficient media (1.6 **·** 10^8^ cells in 50 mL media), the cultures of *M. mazei* ΔsP36 show a significantly prolonged phase of adaptation (38 h lag phase) in the ammonium sufficient medium, before again entering exponential growth phase reaching the same doubling time as the wild type. In contrast, the wild type (wt) is immediately entering exponential growth after the shift to ammonium sufficiency. These findings strongly indicate a crucial role of sP36 under nitrogen up-shift conditions.

**Figure 1:**
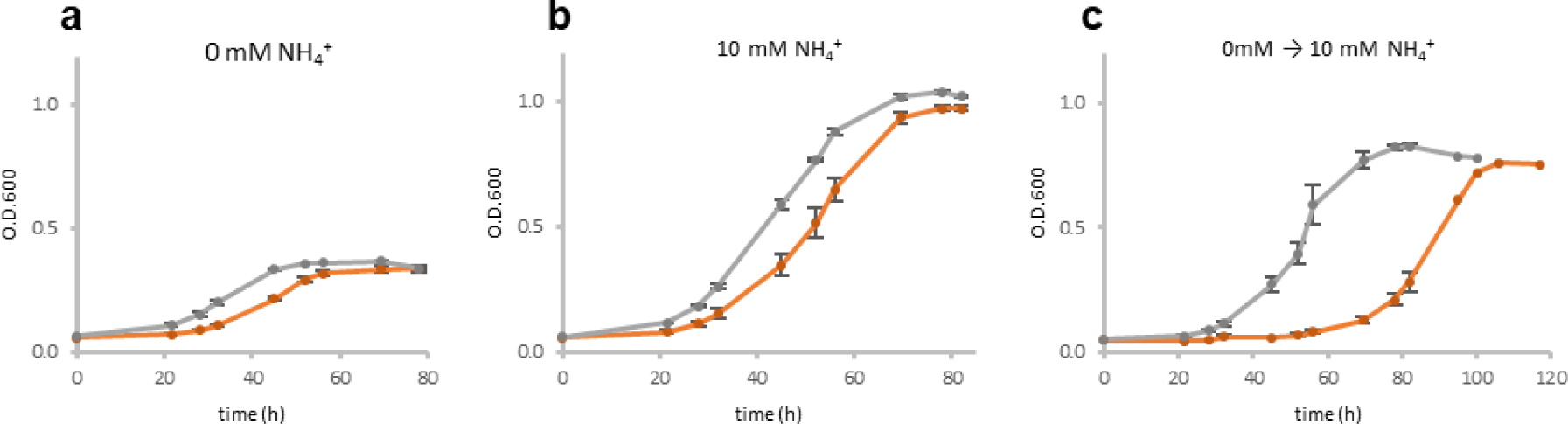
Growth analysis of *M. mazei* ΔsP36 in comparison to *M. mazei* wt: Growth of the *M. mazei* sP36 deletion mutant (ΔsP36), ●(orange), and the wildtype, ●(grey), grown under different nitrogen availabilities with either 0 mM (**A**) or 10 mM (**B**) NH_4_^+^ in the medium, or shifted from 0 mM to 10 mM (**C**). The respective NH_4_+ concentration is depicted. In each case 50 mL anaerobic minimal medium were inoculated with 1.6 **·** 10^8^ cells (T_0_) The standard deviation of three biological replicates are shown.

### sP36 localizes at the cytoplasmic membrane in response to a shift from N limitation to N sufficiency

The cellular localization of sP36, under N limited growth conditions (-N) and after an ammonium up-shift, was evaluated by subcellular fractionation of the cell extract and subsequent Western blot analysis using peptide antibodies directed against sP36. 1000 mL of *M. mazei* cultures were grown under -N. When reaching mid-exponential growth phase (T_600_ = 0.2), 50 % of the cultures were shifted to N sufficiency by supplementing with 10 mM ammonium (final concentration). The remaining 50 % were kept N limited. After further incubation for 30 min, subcellular fractionation was conducted as described in MM, followed by Western blot analysis of the respective cytoplasmic and membrane fractions using specific peptide antibodies against sP36. Overall, three independent biological replicates were analyzed, each with three technical replicates. Under –N, sP36 was predominantly dispersed in the cytoplasm (93 ±3 %). However, upon the ammonium up-shift, most of sP36 relocated into the membrane fraction (57 ±10 %; Fig. 2B).

**Figure 2:**
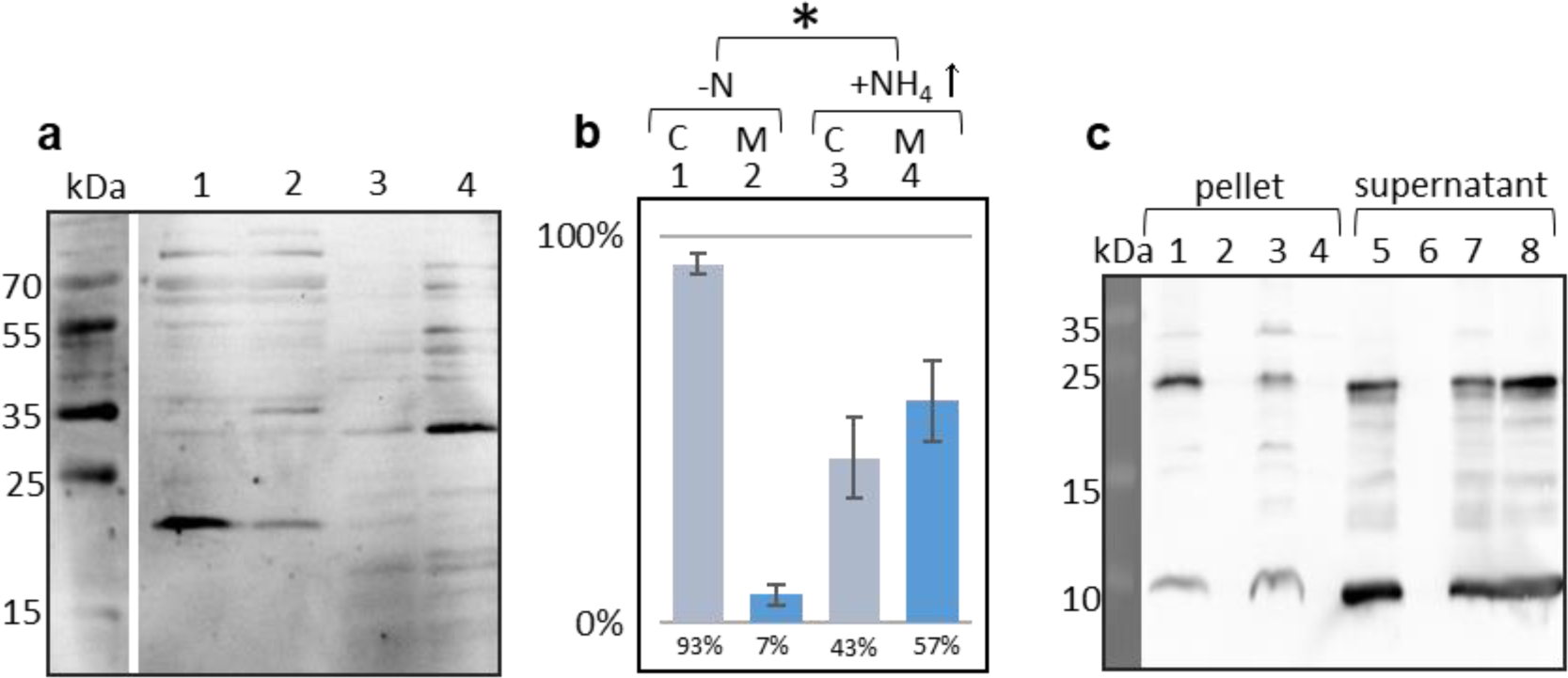
Interaction of sP36 with the cytoplasmic membrane of *M. mazei* under different N conditions. **A+B:** *M. mazei* cultures were grown under N limitation (0 mM NH_4_^+^ N_2_, (**-N**). When T_600_ of 0.2 was reached 50 % of the cultures were shifted to N sufficiency (10 mM NH_4_^+^ final concentration, **NH_4_ ↑**). **A:** Membrane- and the cytoplasmic fractions were analyzed by western blot with polyclonal antibody raised against sP36. **1:** -N cytoplasmic fraction; **2**:, NH_4_↑ cytoplasmic fraction; **3**: -N membrane fraction; **4**:, NH_4_↑ membrane fraction. Depicted is one exemplary Western blot out of three biological replicates. **B: Relative quantification of the dominant bands of sP36 subcellular fractions from *M. mazei.* 1: -**N cytoplasmic fraction; **2: -**N membrane fraction; **3:**, NH_4_↑ cytoplasmic fraction; **4**: NH_4_↑ membrane fraction; the distribution of the sP36 subcellular localization was calculated based on three biological replicates. The amount of sP36 in the cytoplasm and the membrane fraction of one culture was set to 100 %. Significance was tested using two-tailed t-test. *P = 0.014; df = 4. **C: In vitro interaction of sP36 with *M. mazei* ΔsP36 membrane fractions:** Untagged sP36 (100 µg derived from His_6_-SUMO-sP36) was incubated together with the membrane fraction of a *M. mazei* ΔsP36 subcellular fractionation. **Lanes 1-4: pellet of 210,000 g centrifugation; lane 1**: sP36 incubated with –N membrane fraction of *M. mazei* ΔsP36, lane **2**: -N membrane fraction of *M. mazei* ΔsP36 without sP36 (control); **3**: sP36 incubated with membrane fraction of *M. mazei* ΔsP36 grown under N sufficiency (+N); **4**: pellet of sP36 without membrane fraction (control); **lanes 5-8: supernatant of 210,000 g centrifugation**; **5**: sP36 incubated with -N membrane fraction of *M. mazei* ΔsP36, **6**: -N membrane fraction of *M. mazei* ΔsP36 without sP36 (control); **7**: sP36 incubated with membrane fraction of *M. mazei* ΔsP36 grown under N sufficiency (10 mM) ; **8**: sP36 without membrane fraction (control). Depicted is one exemplary Western blot out of three biological replicates.

**Figure 3:**
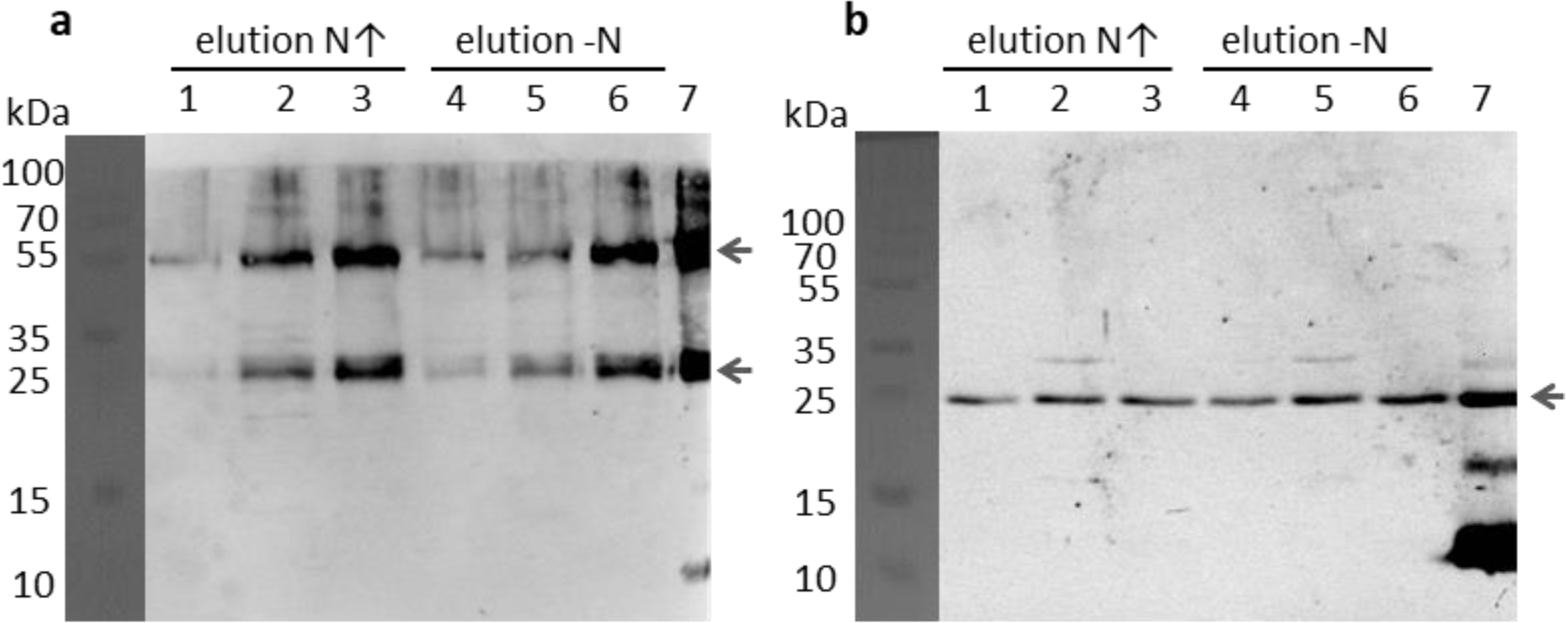
Pulldown with AmtB_1_-His_6_ as bait against *M. mazei* cell extract grown under N limited (–N) and shifted to nitrogen sufficient (N↑) condition. **A:** Western blot using anti-His-tag antibody shows AmtB_1_ in the elution fractions of the N↑ and –N pulldowns. Indicated by a black arrow are the monomeric AmtB_1_ at approximately 30 kDa and the dimeric AmtB_1_ at 55 kDa. **B:** Western blot with specific sP36 peptide antibodies shows coelution of chromosomal expressed native sP36 from *M. mazei* cell extract with AmtB_1_-His_6_. **1-3:** elutions of the pulldown using cell extract of cells grown under N up-shift, **4-6:** elutions of the pulldown using cell extract of cells grown under N limited condition, **7:** positive controls, purified AmtB_1_-His_6_.(A) and purified sP36 (B), respectively.

The observed interaction between sP36 and the cytoplasmic membrane was further verified in an *in vitro* assay using purified tag-less sP36 and cytoplasmic membrane fractions. Membrane fractions from the mutant *M. mazei* strain ΔsP36 grown under -N, as well as after an ammonium upshift (+NH_4_ ^+^↑) were generated by ultracentrifugation as described in MM. The heterologous expressed, purified tag-less sP36 (100 µg) was incubated in the presence of the *M. mazei* ΔsP36 membrane fractions (5 mg) for 15 min at RT followed by ultra-centrifugation at 210,000 *g* and 4°C for 1 h. The respective pellet and supernatant were evaluated for the presence of sP36 by Western blot analysis using peptide antibodies against sP36 (see Fig. 2 C). In the control sample of *M. mazei* ΔsP36 membrane (-N) without prior incubation with sP36, no signal was detected, neither in the supernatant (lane 6), nor in the membrane fraction (pellet, lane 2), confirming the absence of sP36 in the *M. mazei* ΔsP36 mutant strain as well as the specificity of the antibody. In the absence of membrane fractions, purified sP36 was exclusively present in the supernatant after ultracentrifugation (lane 8 vs. lane 4). However, when incubated in the presence of the cytoplasmic membrane fraction of cells grown under nitrogen limitation, approximately 50 % of sP36 was detected in the membrane fraction (lane 1, pellet), strongly arguing for a recruiting of the soluble hydrophilic sP36 to the membrane. Even when cells were grown under nitrogen sufficiency, part of the sP36 was detectable in the membrane fraction after centrifugation (lane 3).

### sP36 interacts with ammonium transport proteins

The subcellular localization experiments as shown above strongly suggest a localization of sP36 to the cytoplasmatic membrane upon an ammonium upshift. A potential interacting partner of sP36 in the cytoplasmic membrane is the ammonium transporter AmtB_1_, which is only expressed under N limitation^38^ and is a key component under N-limitation for transporting residual ammonium into the cell, which critically requires to be inhibited upon an up-shift. To test this hypothesis, a pull-down experiment with purified C-terminal His_6_-tag AmtB_1_ was performed using crude cell extracts of *M. mazei* either grown under N limitation or under N limitation but shifted to ammonium sufficiency (0 → 10 mM) in exponential growth phase for 30 min. After incubating AmtB_1_-His_6_ with 25 mg total cell extract, a Ni-NTA affinity chromatography was performed and the elution fractions analyzed by SDS-PAGE and Western blot using anti-His tag and anti-sP36 antibodies. The results clearly show that exogenous AmtB_1_-His_6_ and native sP36 co-elute independent of the nitrogen conditions in which the cells were grown. These findings strongly suggest that AmtB_1_ is forming a complex with sP36. The direct interaction between sP36 and AmtB_1_ was further confirmed and evaluated by microscale thermophoresis (MST) using recombinant AmtB_1_-His_6_ and untagged sP36 (RED) resulting in an estimated dissociation constant of *K*_D_ = 0.26 ± 0.07 µM (Fig. 4A).

**Figure 4:**
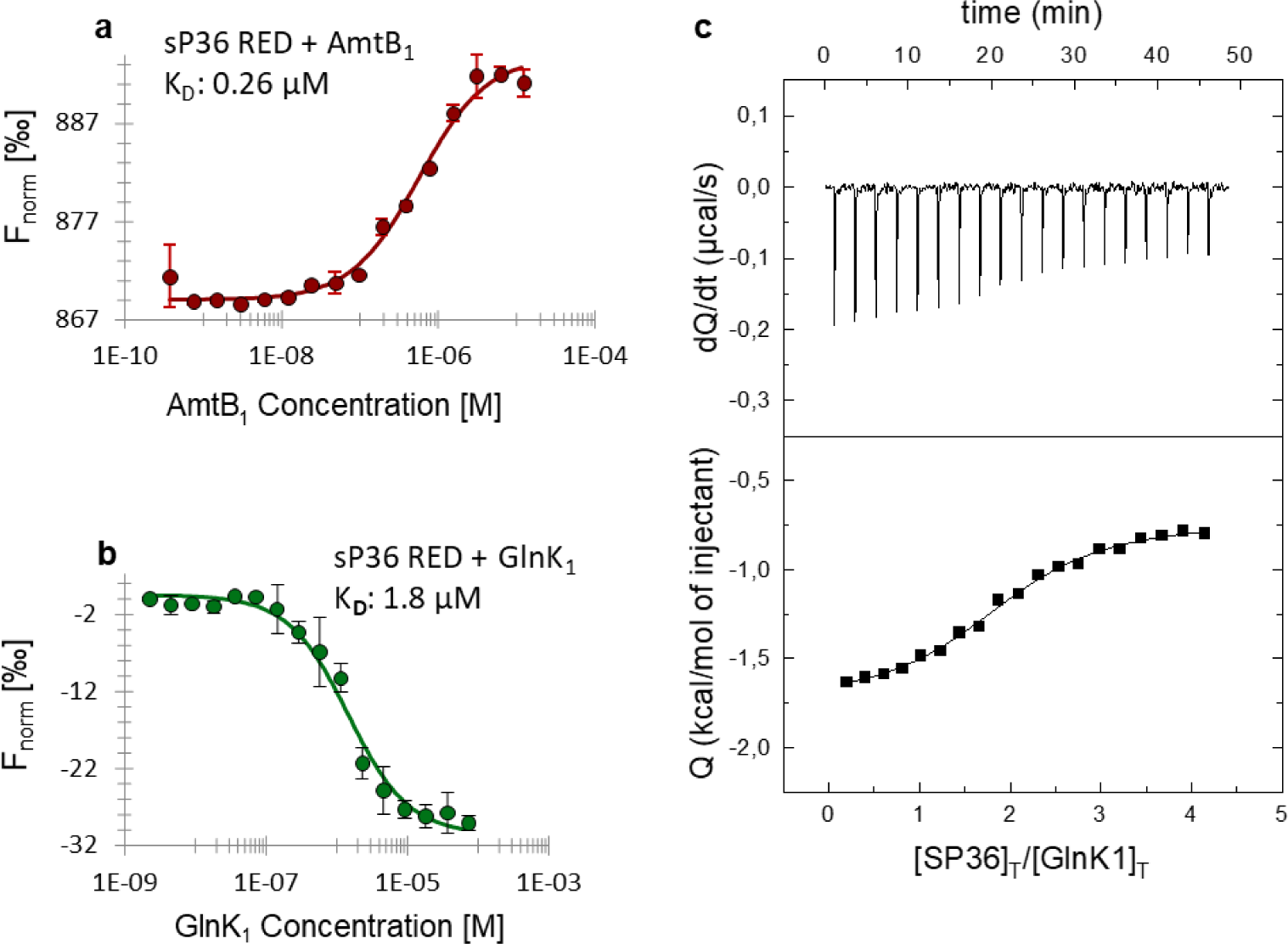
sP36 interaction studies using microscale thermophoresis (MST) and isothermal titration calorimetry (ITC). A: Interaction studies between sP36 and AmtB_1_-His_6_ using MST. RED labeled untagged sP36 was used at 100 nM and AmtB_1_-His_6_ at different concentrations ranging from 12.5 µM to 0.38 nM, calculated based on the monomeric molecular mass. Based on 3 biological replicates the *K*_D_ was estimated to be 0.26 µM (± 0.07 µM). **B: Interaction studies between sP36 and GlnK_1_ using MST.** RED labeled untagged sP36 at 100 nM and His_6_-GlnK_1_ in 16 different concentrations ranging from 7.5 µM to 0.23 nM were used for MST analysis, resulting in a dissociation constant *K*_D_ of 1.8 µM (± 1.1 µM). In both cases (A and B), exemplarily one of three biological replicates is depicted. **C**: **Interaction studies between sP36 and GlnK_1_ using ITC.** 20 μM His_6_-GlnK_1_ was titrated with 300 μM at 25 °C. Control experiments were performed by injecting sP36 into buffer. The dissociation constant *K*_D_ was evaluated to be 5.4 µM. The sP36 - GlnK_1_ interaction has a stoichiometry of 2:1 as described in MM.

As stated in the Introduction, in bacteria, the PII-like protein GlnK is known to interact with the ammonium transporter AmtB to modulate the transport activity of AmtB in response to an ammonium upshift^49^. Therefore, we next aimed to evaluate the interaction between sP36 and GlnK_1_, by using recombinant N-terminal his-tagged GlnK_1_ (His_6_-GlnK_1_) and sP36 (RED) proteins by MST, which yielded in a dissociation constant of 1.8 ± 1.1 µM (Fig. 4B).

The interaction between sP36 and GlnK_1_ was further verified by isothermal titration calorimetry (ITC, Fig. 4C). The ITC analysis clearly verified the interaction and demonstrated that sP36 forms a complex with GlnK_1_ with a dissociation constant *K*_D_ of 5 µM in a 2:1 stoichiometry, *i.e.*, each monomer of GlnK_1_ binds two sP36 monomers.

Given that GlnK_1_ forms trimers in solution^27^, a 2:1 stoichiometry aligns perfectly with the oligomeric state of sP36, which has been determined to form stable hexamers in solution by analytical size exclusion chromatography (SEC) experiments. SEC was performed with sP36 from heterologous expression in *E. coli* as well as with His_6_-sP36 purified from *M. mazei* (Fig. 5A and 5B). Using AlphaFold2 for computational modeling, we generated a structural model of hexameric sP36 (Fig. 5C), whose statistical parameters pLDDT and PAE (Supplemental Figure A1) demonstrate that the residues are correctly positioned in their local environment, and the monomers are accurately positioned relative to each other^50^. The hexameric configuration of sP36 adopts a truncated cone shape, with negative charged surface at the narrower side (Fig. 5D).

**Figure 5:**
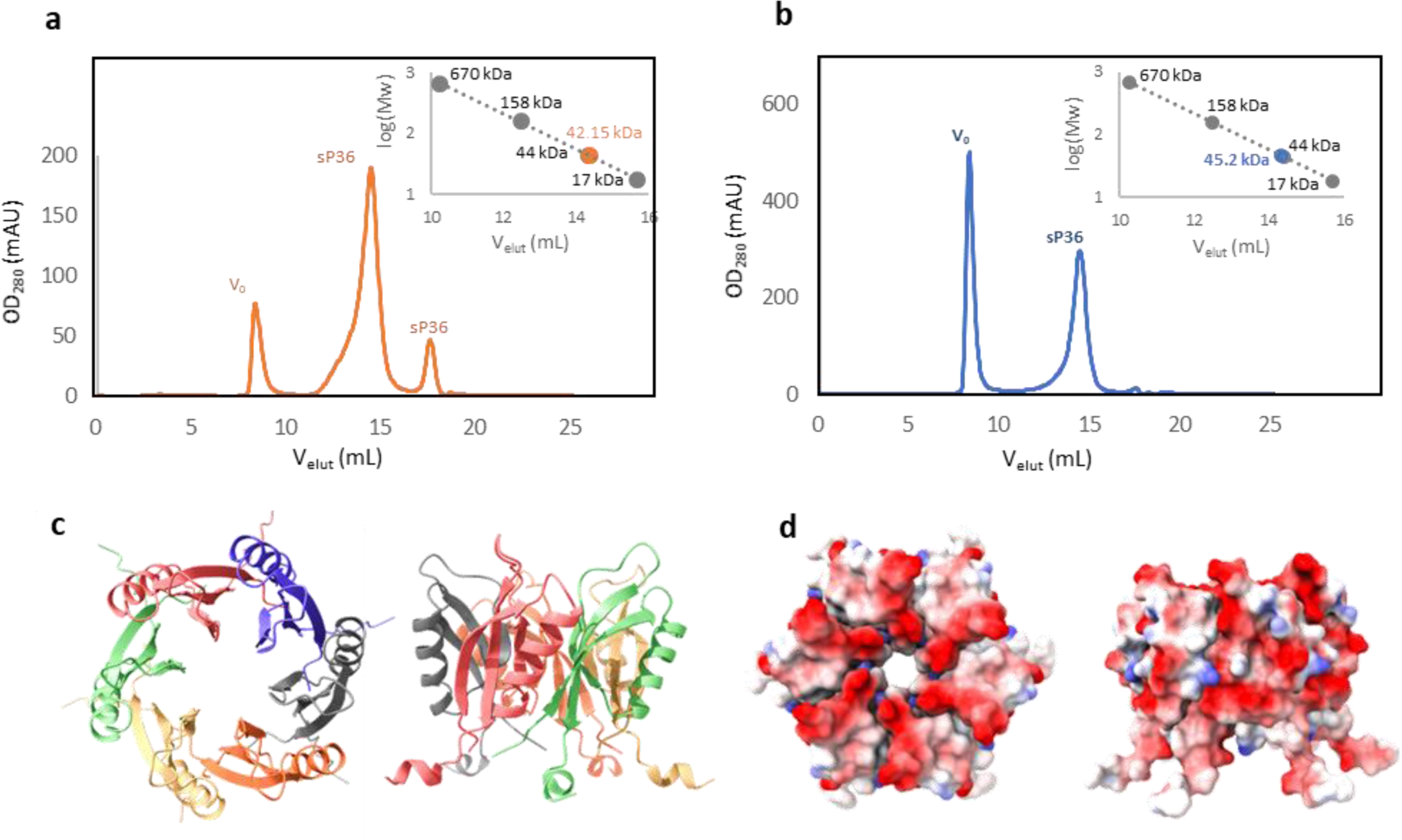
Oligomerization studies of purified sP36. **A:** Untagged sP36 from expression in *E. coli*. Proteins and protein complexes were separated in a size exclusion chromatography. The fractions of the main elution peak (retention volume of 14 mL to 15 mL) correspond to the molecular mass of sP36 hexamer. ●(orange): sP36-His_6_, ●(grey): size exclusion standard. **B:** His_6_-sP36 from expression in *M. mazei*. Proteins and protein complexes were separated in a size exclusion chromatography. The fractions of the main elution peak (retention volume of 14 mL to 15 mL) correspond to the molecular mass of His_6_-sP36 hexamer. ●(blue): sP36-His_6_, ●(grey): size exclusion standard. **C + D:** Structure prediction of a sP36 hexamer ^50,58^. **C:** Secondary structural elements of sP36 hexamer, each protomer in a different color. **D:** Electrostatic surface of the sP36 hexamer. The calculation was performed with APBS plug-in implemented in PyMOL (Schrödinger Inc. (2015) The PyMOL Molecular Graphics System. Version 2.0 Schrödinger LLC). Color oscillates from −2.0 (red) to +2.0 (blue) KbT/ec.

MST and the ITC independently showed an interaction between sP36 and GlnK_1_ with a high affinity (*K*_D_ in the low µM range). Thus, we next aimed at evaluating the impact of sP36 on the subcellular localization of GlnK_1_. *M. mazei* wt and ΔsP36 strains were grown under N limitation followed by an ammonium upshift for 30 min and cultures harvested in mid exponential growth phase. After subcellular fractionation, the presence of GlnK_1_ was detected in the cytoplasmic and membrane fractions by Western blot. In the wt strain, GlnK_1_ was predominantly present in the membrane fraction, while only a minor fraction was detected in the cytoplasm (Fig. 6). However, in ΔsP36, GlnK_1_ is no longer detected in the membrane fraction, but mostly in the cytoplasm. Remarkably, the total amount of GlnK_1_ in the absence of sP36 appears to be decreased (Fig. 6).

**Figure 6:**
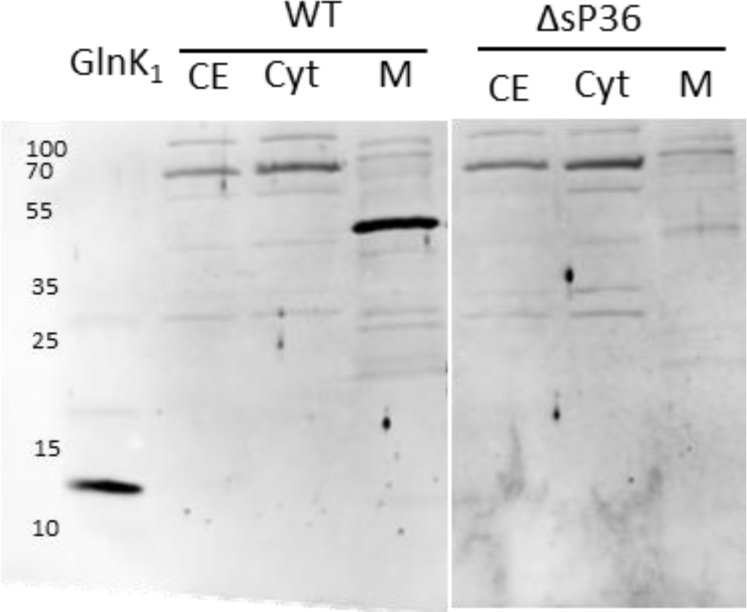
Localization of GlnK_1_ in *M. mazei* ΔsP36. *M. mazei* wt and ΔsP36 cultures (50 mL) were grown under N limitation until reaching exponential growth phase and then shifted to N sufficiency (10 mM). Subcellular fractions were generated. The cytoplasmic and membrane fraction were evaluated for GlnK_1_ presence by Western blot analysis using polyclonal antibodies against GlnK_1_. Cytoplasmic (C) and membrane (M) fraction from *M. mazei* wt(DSM3647) and ΔsP36 were analyzed compared to purified His_6_-GlnK_1_. Depicted is one exemplary Western blot out of three biological replicates.

## Discussion

Based on the aforementioned results, we propose that in *M. mazei* both, GlnK_1_ and sP36, are required for the complete inhibition of AmtB1, the ammonium transporter, in response to an up-shift in ammonium concentration after a period of N limitation in *M. mazei*. The essential role of sP36 in AmtB_1_ regulation during an N-upshift is further corroborated by our genetic analysis. The chromosomal mutant strain (*M. mazei* ΔsP36) displays a significantly prolonged lag phase when shifted from N limitation to ammonium sufficiency (10 mM NH_4_ ^+^) compared to the wild type strain. This strongly indicates that the ammonium transporter in the absence of sP36 retains significant activity, leading to an unnecessary cycle of active AmtB_1_-mediated import together with passive diffusion, resulting in excessive energy consumption. This might explain the significant prolonged lag phase observed in the deletion mutant strain after a shift to N sufficiency.

The mechanism of AmtB regulation by the PII protein GlnK is well characterized in *E. coli*. Here, the cellular nitrogen status is perceived by GlnD, which is transducing the signal to GlnK through a covalent modification. Under N limitation GlnK is uridylylated at the Y51 residue in the T-loop by GlnD. In response to increased ammonium concentrations, GlnK is rapidly deuridylylated by GlnD and the demodified GlnK subsequently interacts with AmtB and inhibits its activity^46,49^. The trimeric GlnK binds AmtB with the T-loop of each monomer physically blocking the hydrophobic pore of the AmtB trimer and the cytoplasmic pore exit^51^. This regulatory mechanism appears to be highly conserved and has been also shown for *Rhodospirillum rubrum*^52^, and the archaea *Haloferax mediterranei*^53,54^ and *Archaeoglobus fulgidus*^55^. The *A. fulgidus* regulation was mainly proposed based on structural studies of purified AmtB and by using a docking model for the interaction with the PII-like protein. Interestingly, the conserved T-loop of the *A. fulgidus* PII-like protein lacks the Y51 residue, which is the residue modified by uridylyation in *E. coli*^55^.

In *M. mazei* two copies of the *glnK*/*amtB* operon are present. While the *glnK_1_*/*amtB_1_* operon is highly regulated in response to N availability by NrpR, the *amtB_2_*/*glnK_2_* operon is not expressed under N limitation and has been proposed to be a potential backup system^38,56^. Although the T-loop of GlnK_1_ contains the conserved tyrosine residue (Y51), no posttranslational modification of GlnK_1_ could be observed in response to an ammonium up-shift in *in vivo* and *in vitro* experiments^27^. Moreover, no GlnD homolog is encoded in the *M. mazei* genome^57^. Consequently, in *M. mazei*, a potential GlnK_1_ regulation of AmtB_1_ requires a different mode of signal perception of changing nitrogen conditions.

Based on our results, we propose a hypothetical model for the regulation of AmtB_1_ by sP36 in response to an increase in N availability, which is summarized in Figure 7. In the absence of combined nitrogen or when the ammonium concentration is very low, most of the sP36 is located in the cytoplasm, while AmtB_1_ actively transports the remaining NH_4_ ^+^ into the cell. When the external ammonium concentration increases (N-up-shift), NH_3_ diffusion provides the cell with sufficient ammonium. Thus, the energy-consuming NH_4_ ^+^ transport by AmtB1 is inhibited through a direct protein-protein interaction with the GlnK_1_ trimer. This complex formation between trimeric AmtB_1_ and trimeric GlnK_1_, however is crucially dependent on hexameric sP36, which binds both proteins with high affinity. Consequently, sP36 favors the AmtB_1_-GlnK_1_ interaction in response to an ammonium up-shift after a period of N limitation. The structure of the binary and ternary complexes and the hierarchy of the interactions requires further investigation and is currently being studied in our laboratory. We also note that the negatively charged surface on sP36 (Fig. 5D) is well-suited to interact with the positively charged intracellular side of AmtB_1_, as reported in the literature.

**Figure 7:**
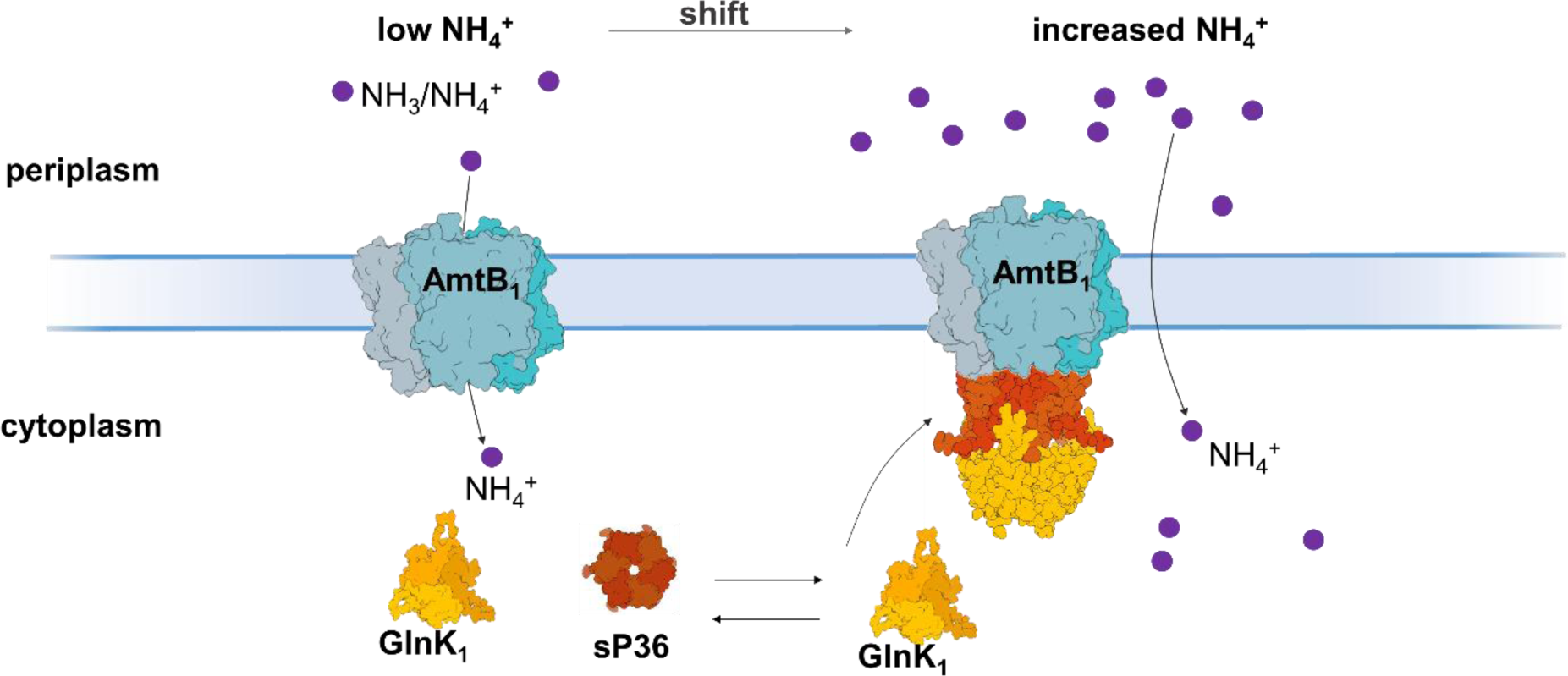
Hypothetical model of AmtB_1_ regulation in *M. mazei*. The AmtB_1_ trimer is actively importing ammonium under low N concentration. However, after an up-shift to ammonium sufficiency in the surrounding medium, sP36 is proposed to mediate the formation of a GlnK_1_-AmtB_1_ complex, allowing the GlnK_1_ trimer to block AmtB_1_ activity to completeness. Displayed structures of GlnK1 and sP36 were generated using AlphaFold2 ^57,58^ (supplemental Figure A2). The structure of the ternary complex is a hypothetical model and requires further investigation.

Finally, we speculate that sP36-mediated inhibition of AmtB_1_ represents an ancient mechanism for responding to changes in N availability, predating the AmtB inhibition by a PII-like protein before the GlnD-dependent uridylylation was invented.

## Methods

### Construction of plasmids

#### sP36 genomic deletion

The flanking regions ∼1,000 bp upstream and ∼1,000 bp downstream of the sORF36 were amplified from genomic *M. mazei* DNA by using primers (Eurofins, Ebersberg, Germany) listed in table A2. A puromycin resistance (Pur^R^) mediating *pac-*cassette was restricted from pRS207. The 1,000 bp downstream fragment was restricted by enzymes *EcoRI* and *BamHI* and inserted into the multiple cloning site (MCS) of the vector pMCL210, resulting in pRS1305. The 1,000 bp upstream fragment was cleaved using *EcoRI* and *KpnI* and subsequently ligated in pRS1305. The resulting plasmid was designated pRS1307. The *pac*-cassette was ligated into the *EcoRI* site of pRS1307 and the resulting plasmid was named pRS1308. pRS1308 was linearized using *ScaI* and transformed into *M. mazei* wt (3A) cells by liposome mediated transformation. Insertion in the chromosome occurred through double homologous recombination by selection for puromycin resistance^56^. The success of the allelic marker exchange of single mutant colonies was verified in puromycin containing media and further analyzed via southern blot analysis with specific probes directed against the *sP36* gene and the *pac*-cassette (supplemental Figure A3).

#### sP36 overexpression

For cloning *MMsORF36* into pETSUMO, pET28a and pWM321 a construct including the *pmcrB* promotor and a (His)_6_-*MMsORF36* fusion was synthesized (Eurofins Genomics, Ebersberg, Germany) The plasmid was named pRS1214. *MMsORF36* was cloned into pETSUMO using pRS1214 as template, primer pair sORF36_3for/sORF36_3rev and the Champion™ pET SUMO Expression System (Thermo Fisher Scientific, Waltham, USA) according to the manufactures instructions, yielding plasmid pRS1240 and strain *E. coli* BL21 K4099. The cloning of *MMsORF36* into pET28a was done via an intermediate. First, *MMsORF36* was amplified with the primer pair sORF36_forNdeI/sORF36_3revNdeI (table A2), using the template pRS1214 and subsequently ligated via TA cloning into the pCRII vector (Thermo Fisher Scientific, Waltham, USA) according to the manufacturer instructions. The resulting construct was designated pRS1223 in *E. coli* DH5α K4071. In the second step, the insert was excised from pRS1223 using the *NdeI* site and ligated into the *NdeI* site of pET28a. The resulting plasmid was designated pRS1225 and transformed in *E. coli* BL21 pRIL yielding the strain K4092. Cloning of *MMsORF36* in pWM321 the *pmcrB* promotor-(His)_6_-*MMsORF36* fusion was isolated using *SacI* and *KpnI* sites from pRS1214 and ligated into the corresponding sites of pWM321, resulting in plasmid pRS1227. pRS1227 was transformed into *M. mazei* wt (3A) as described before^56^.

#### His_5_-SUMO-TEV-sP36 construct

A TEV cleavage site was inserted between the Sumo tag and sORF36 in the His_5_-Sumo-sP36 overexpression construct (pRS1240). Therefore, site directed mutagenesis was conducted using primers SP36_TEV_rv and SP36_TEV_fw (table A2).

#### Growth of M. mazei

*M. mazei* was cultivated in sealed bottles of anaerobic minimal medium with a gaseous phase consisting of N_2_ and CO_2_ (vol/vol, 80/20)^26,57^. The medium was supplemented with 150 mM methanol as carbon and in case of cultures growing in nitrogen sufficiency, additionally with 10 mM ammonium chloride. Cells were cultivated until an optical turbidity of 0.5-0.6 at 600 nm (T600 = 0.5-0.6). –N Cells were grown until T600 = 0.2-0.3.

*M. mazei* cells were harvested by centrifugation at 4,000 g at 4°C for 30 min. The cells were resuspended in 2 mL 50 mM Tris buffer (pH 7.6) and lysed by using a dismembrator (Sartorius, Göttingen, Germany) at 1,600 rpm for 3 min. The whole cell extract was centrifuged for 30 min at 13,000 g and 4 °C to get rid of cell debris and remaining unlysed cells.

### Subcellular fractionation

For subcellular fractionation of *M. mazei*, the cultures were grown anaerobically as described. Cells were harvested by centrifugation with 6,000 g at 4°C for 30 min. The cells were resuspended in 10 mL Tris buffer (50mM, pH 7.6) and afterwards lysed by using a dismembrator (Sartorius, Göttingen, Germany) at 1,600 rpm for 3 min. The lysate was centrifuged for 30 min at 7,500 g and 4°C. To separate membrane and cytoplasmic fractions, the supernatant was further centrifuged at 210,000 g for 1 h at 4°C.

### Purification of expressed proteins

#### His_6_-sP36, His_6_-GlnK1, His_6_-SUMO-sP36

*E. coli* Bl21 (DE3)/pRIL cultures were grown in LB-medium at 37°C under continuous shaking. At T600 = 0.6, protein expression was induced by adding 100 µM IPTG (final concentration) and the cultures were incubated for two hours. Cells were harvested by centrifugation at 4,000 g at 4°C for 20 min, suspended in 6 mL phosphate buffer A (50 mM phosphate, 300 mM NaCl, pH 8.0) and lysed by passing the French pressure cell two times with 80 N(mm^2^)^-1^. Afterwards the extract was centrifuged for 30 min at 13,000 g and 4 °C to get rid of cell debris and remaining unlysed cells. For protein purification from *M. mazei*, cultures were grown and lysed as described above. His-tagged proteins were purified by affinity chromatography on Ni-NTA agarose (Qigen, Hilden, Germany) gravity flow columns with 1 mL bed volume. Proteins were eluted in steps with 100 mM, 250 mM and 500 mM imidazole.

### SUMO cleavage

For cleavage of the SUMO-(His)_6_-tag, 200 µL of SUMO-protease (Thermo Fisher Scientific, Waltham, MA, USA) per 1 mg tagged protein was used. The reaction was incubated for 1 h at 30 °C. The cleaved sP36 protein was afterwards purified by a second step of affinity using Ni-NTA agarose (Qiagen, Hilden, Germany) in phosphate buffer A.

#### His_5_-SUMO-TEV-sP36 construct

*E. coli* Rosetta containing the His_5_-SUMO-TEV-sP36 construct was grown in LB-medium at 37°C under continuous shaking. At T600 = 0.6, protein expression was induced by adding 100 µM IPTG (final concentration) and the cultures were incubated over night at 20°C. Cells were harvested by centrifugation at 4,000 g at 4°C for 20 min, suspended in Tris-HCl buffer A (20 mM Tris-HCl, 300 mM NaCl, pH 8.0) and lysed by sonication (Gardiner, NY, USA). Afterwards the extract was centrifuged for 30 min at 13,000 g and 4 °C. His-tagged proteins were purified by affinity chromatography in a His-Trap column (Cytiva, Marlborough, MA, USA). Proteins were eluted with Tris-HCl buffer with 500 mM imidazole. Protein fractions were dialyzed against 20 mM Tris-HCl, 0.5 M NaCl, 2 mM DTT, pH 8.0 in the presence of TEV-protease. The cleaved sP36 protein was afterwards separated from undigested tagged sP36, cleaved His_5_-SUMO-TEV-tag as well as the His-tagged TEV protease by a second step of affinity using a His-Trap column (Cytiva, Marlborough, MA, USA). Final purification was conducted using gel filtration with a S-100 Sephacryl HR column (Cytiva, Marlborough, MA, USA) and 20 mM Tris-HCl, pH 8.0, 0.15 M NaCl buffer.

#### AmtB_1_-His_6_

AmtB_1_-His_6_ was purified from heterologous overexpression in *E. coli* C43 using solubilized membrane fraction. Therefore, cultures were grown in LB-medium at 37°C. At T600 = 0.6, protein expression was induced by adding 500 µM IPTG (final concentration) and the cultures were incubated for three hours at 37°C. Cells were harvest by centrifugation at 6,000 g, for 20 min at 4 °C. 4 g pellet was resuspended in 4 mL 50 mM Tris buffer (pH7.6) and lysed by passing through the French press two times at 40 N(mm^2^)^-1^. The extract was then centrifuged again at 8,000 g for 20 min at 4°C to remove remaining unlysed cells and cell debris. The cleared supernatant was transferred into new tubes and centrifuged in an ultracentrifuge (Optima XPN-100 Ultracentrifuge, Beckmann coulter, Brea, California, USA) for 1 h at 210,000 g and 4 °C. The membrane pellet was washed with 15 mL 50 mM Tris buffer (pH 7.6) and again centrifuged in the ultracentrifuge for 1 h at 4 °C and 210,000 g. Afterwards the membrane proteins in the pellet were solubilized in 1 mL of phosphate buffer B (50 mM phosphate, 150 mM NaCl, 2% DDM, pH 8.0).

The solubilized membrane fraction was added on an affinity chromatography Co-NTA agarose gravity flow column (bed volume: 0.5 mL). For washing and elution steps, phosphate buffer C (50 mM phosphate, 150 mM NaCl, 0.05 %DDM, pH 8.0) was used. His-tagged proteins were eluted in 0.5 mL steps using phosphate buffer C with 100 mM, 250 mM and 500 mM imidazole.

### Microscale thermophoresis (MST)

Proteins were purified to apparent homogeneity by affinity chromatography using Ni-NTA agarose and labeled with the RED-NHS, 2^nd^ generation, 650 nm fluorescent dye using the Monolith NT RED-NHS lysine labeling kit according to manufacturer’s protocol (NanoTemper, Munich, Germany). RED labeled untagged sP36 at 100 nM and His6-GlnK1 in 16 different concentrations ranging from 7.5 µM to 0.23 nM or 10 nM of sP36-RED and AmtB_1_-His_6_ in 16 dilutions ranging from 12.6 µM to 0.38 nM (all concentrations based on monomeric molecular mass) was used. The protein interaction was measured in standard capillaries (NanoTemper), 100% excitation power and medium or high MST power (IR-laser intensity). Both interactions were tested in three biological replicates.

### Isothermal titration calorimetry (ITC)

Standard ITC experiments were performed using an Auto-iTC200 system (MicroCal, Malvern-Panalytical, Malvern, UK). Briefly, 20 μM GlnK1 was titrated with 300 μM sP36 in buffer 100 mM potassium phosphate, 2 mM EDTA, pH 7.0 at 25 °C. Control experiments were performed by injecting the sP36 protein into the buffer. The heats of dilution were negligible. The resulting heats were integrated and normalized by ligand injected, and fitted with a model for a single ligand binding site implemented in the software package Origin 7.0 (OriginLab Corporation, Northamptom, MA, USA) employing user-defined fitting routines.

### Size exclusion chromatography

Size exclusion chromatography (SEC) was conducted with 0.3 mg His_6_-sP36, purified from homologous expression in *M. mazei* and with 0.5 mg untagged sP36 (derived from His_6_-SUMO-sP36) using 50 mM phosphate buffer containing 300 mM NaCl, and the analytical gel filtration column ENrich TM SEC 650 (BioRad, Hercules, USA). The protein was eluted with a flow rate of 1 mL min^-1^, with 50 mM phosphate buffer (150 mM NaCl, pH 8.0). Elutions were collected in 1 mL fractions. In order to calibrate the chromatograph, a protein mix (Bio-Rad size exclusion standard; #151-1901, Bio-Rad, Hercules, USA) was used as a standard.

## Supporting information

Supplemental Table 1: Strains and plasmids

Supplemental Table 2: Oligonucleotidess

Supplemental Figures: AlphaFold predictions

Supplemental Figure: sP36 deletion southernblot

## Appendix

**Table A1:** Used Strains and plasmids.

| Archaea |  |  |
| --- | --- | --- |
| ***Methanosarcina mazei* Goe1** | Wildtype | German Collection of Microorganisms and Cell Cultures GmbH (DSMZ) number 3647 |
| ***Methanosarcina mazei* wt (3A)** | Wildtype, improved growth on solid media | <sup>56</sup> |
| ***Methanosarcina mazei* ΔsP36** | *M. mazei* 3A/ΔsP36 genomic deletion | This work |
| ***M. mazei* 3A pWM321/MMsORF36.N** | pRS1227 in *M. mazei* 3A Pur<sup>R</sup> | This work |
| Bacteria |  |  |
| ***E. coli* DH5α** | host strain for cloning of plasmid constructs (F<sup>-</sup>, *lacZ*D<sup>M</sup>15, *recA*1, *hsdR*17, *supE*44, *lacZ*YA, *argF*) | <sup>59</sup> |
| ***E. coli* BL21 (DE3) containing pRIL** | strain for overexpression of archaeal proteins F<sup>-</sup>, *ompT* *hsdS*(rB - mB), *dcm*+, *Tet*r *gal* λ (DE3), *endA* *Hte* [*argU* *proL* *Cam*r] [*argU* *ileY**leuW* *Strep*/*Spec*r] | Stratagene, La Jolla, USA |
| ***E. coli* C43 (DE3)** |  | <sup>60</sup> |
| **E2407** | pWM321 in *E. coli* JM109 λ pir Amp<sup>R</sup>, Pur<sup>R</sup> | <sup>56</sup> |
| **E1570** | pET28a(+) in *E. coli* DH5α Kan<sup>R</sup> | Novagen |
| **K4057** | pRS1214 in *E. coli* DH5α Amp<sup>R</sup> | This work |
| **K4071** | pRS1223 in *E. coli* DH5α Amp<sup>R</sup>, Kan<sup>R</sup> | This work |
| **K4076** | pRS1225 in *E. coli* DH5α Kan<sup>R</sup> | This work |
| **K4078** | pRS1227 in *E. coli* JM109 λ pir Amp<sup>R</sup>, Pur<sup>R</sup> | This work |
| **K4098** | pRS1240 in *E. coli* DH5α Kan<sup>R</sup> | This work |
| **K4099** | pRS1240 in *E. coli* BL21 Kan<sup>R</sup> | This work |
| Plasmids |  |  |
| **pmcl210** | cloning Vector, P15Aori, *lacZ*, *cat* | <sup>61</sup> |
| **pRS207** | pac cassette in pBlueskript SK | <sup>56</sup> |
| **champion pET SUMO** | expression vector, Kan<sup>R</sup> | Thermo Fischer Scientific, Waltham, USA |
| **pWM321** | shuttle Vector *M. mazei* /*E. coli* ori<sup>R6K</sup>: pC2A replicon, Amp<sup>R</sup>, Kan<sup>R</sup> | <sup>62</sup> |
| **pET28a(+)** | expression vector, Kan<sup>R</sup> | Novagen, Darmstadt, Germany |
| **pET21a(+)** | expression vector, Amp<sup>R</sup> | Novagen, Darmstadt, Germany |
| **pCR<sup>TM</sup>II-TOPO®** | Plac, LacZα-ccB, Amp<sup>R</sup>, Kan<sup>R</sup>, pUC ori | Thermo Fisher Scientific, Waltham, USA |
| **pRS1223** | *pCR<sup>TM</sup>II-TOPO®/MMsORF36 Amp<sup>R</sup>, Kan<sup>R</sup>* | This work |
| **pRS1225** | *pET28a/MMsORF36 Kan<sup>R</sup>* | This work |
| **pRS1227** | *pWM321/MMsORF36.N Pur<sup>R</sup>, Amp<sup>R</sup>* | This work |
| **pRS1240** | *pETSUMO/MMsORF36.N Kan<sup>R</sup>* | This work |
| **pRS1308** | pWM321/MMsORF36 flanking regions/pac cassette | This work |
| **pRS1859** | pET21a/MM733 Amp<sup>R</sup> | This work |

**Table A2:**
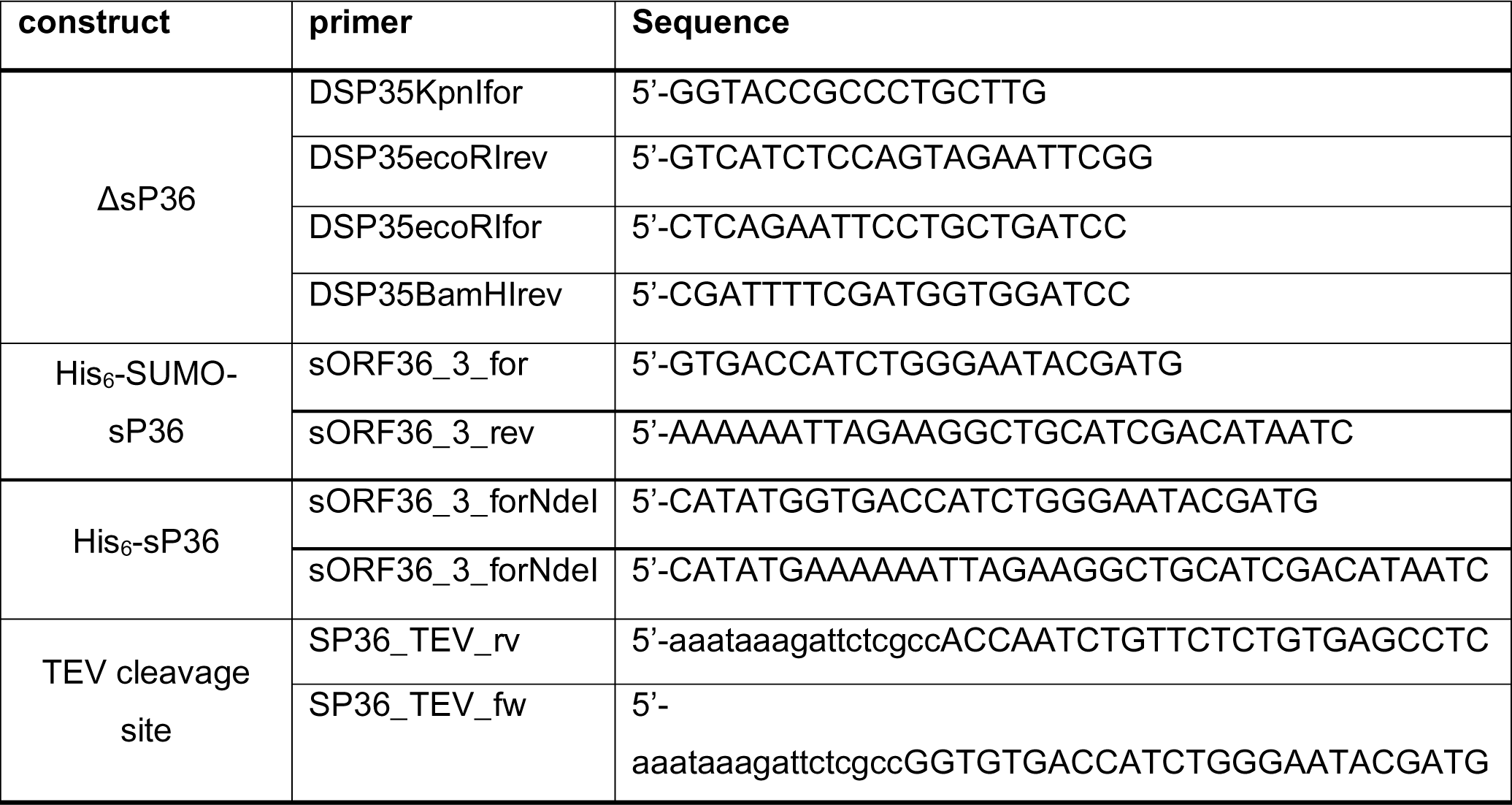
Used Oligonucleotides.

## Supplemental figures

**Supplemental figure A1.**
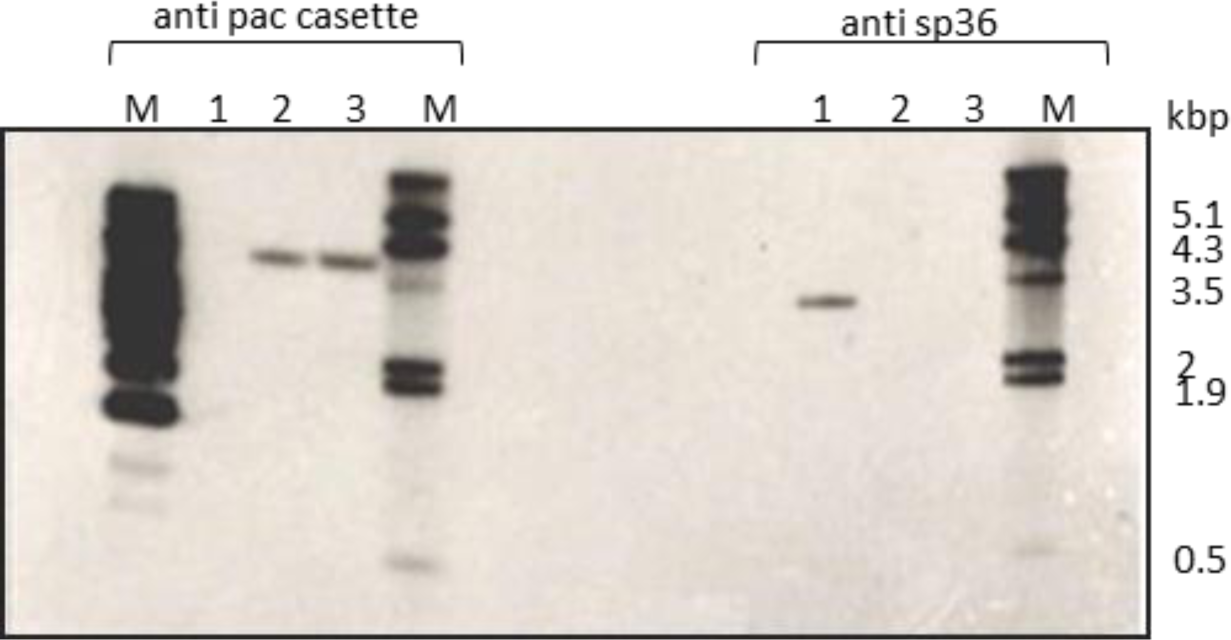
Southern blot of *M. mazei* Δsp36 genomic DNA: to test the success of the sp36 deletion in *M. mazei,* genomic DNA of the new Δsp36 mutant and the wildtype (control) was cleaved by restriction enzyme HindIII and was analyzed with specific probes against sp36 and against the pac casette. **1**: *M. mazei* 3A (wt), **2**: *M. mazei* Δsp36 (clone1), **3**: *M. mazei* Δsp36 (clone2).

**Supplemental figure A2.**
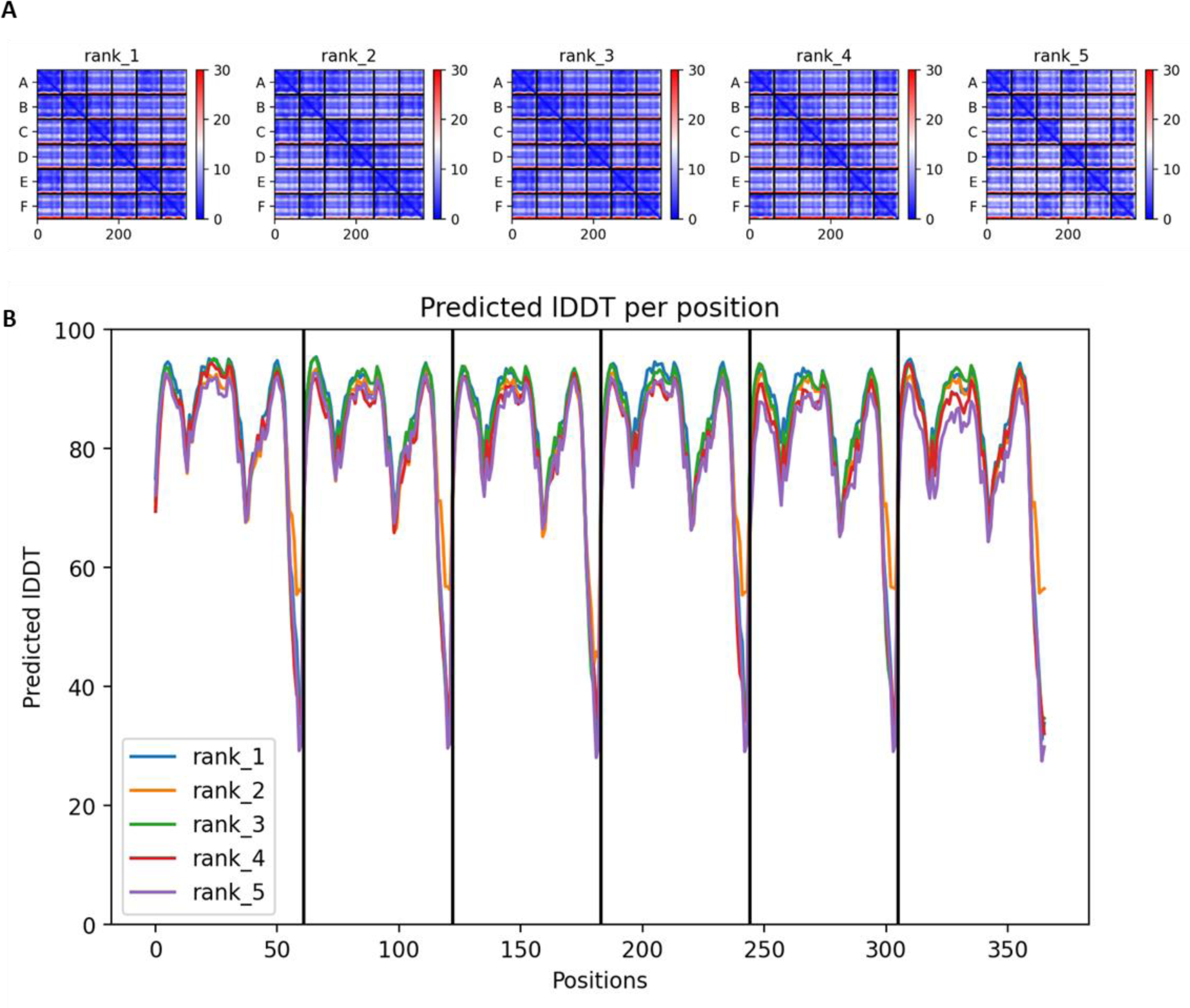
Confidence of the predicted structure of sP36. (A) Predicted aligned error (PAE) and (B) Predicted local distance difference test (IDDT) for the five models generated by AlphaFold2 ^50,58^.

**Supplemental figure A3.**
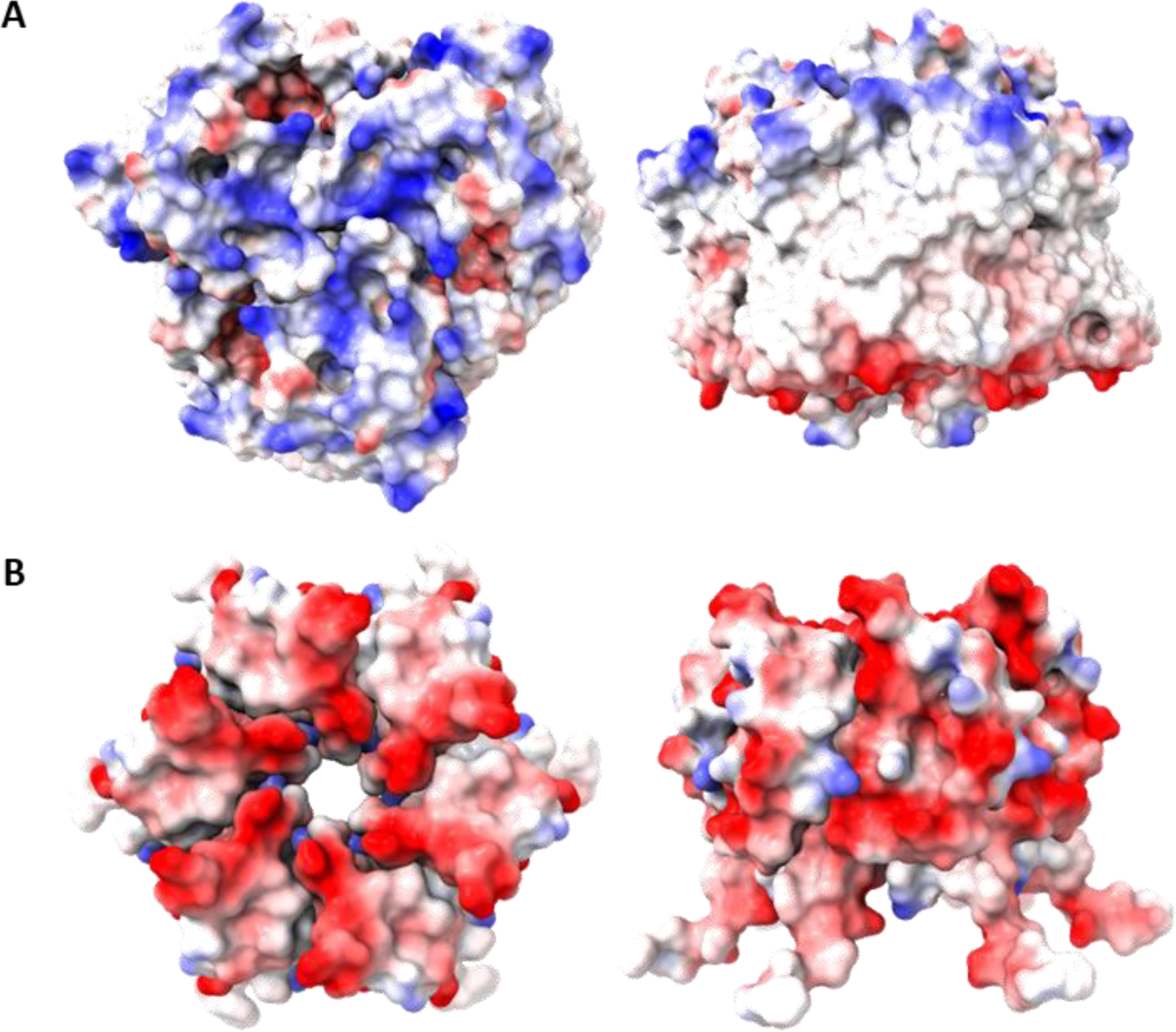
Color-coded representation of the electrostatic potential of the surface of (A) AmtB1 and (B) sP36. The calculation was performed with APBS plug-in implemented in PyMOL (Schrödinger Inc. (2015) The PyMOL Molecular Graphics System. Version 2.0 Schrödinger LLC). Color oscillates from −2.0 (red) to +2.0 (blue) KbT/ec.

