## Supplemental Table 1: Strains and plasmids for "Small protein mediates inhibition of ammonium transport in Methanosarcina mazei – an ancient mechanism?"

### Table S1: Used Strains and plasmids

| | | |
|---|---|---|
| **Archaea** | | |
| ***Methanosarcina mazei* Goe1** | Wildtype | German Collection of Microorganisms and Cell Cultures GmbH (DSMZ) number 3647 |
| ***Methanosarcina mazei* wt (3A)** | Wildtype,<br>improved growth on solid media | (57) |
| ***Methanosarcina mazei* ΔsP36** | *M. mazei* 3A/ΔsP36 genomic deletion | This work |
| ***M. mazei* 3A pWM321/MMsORF36.N** | pRS1227 in *M. mazei* 3A Pur$^{R}$ | This work |
| **Bacteria** | | |
| ***E.* coli DH5α** | host strain for cloning of plasmid constructs (F-, *lacZDM15*, *recA1*, *hsdR17*, *supE44*, *lacZYA*, *argF*) | (59) |
| ***E.* coli BL21 (DE3) contianing pRIL** | strain for overexpression of archaeal proteins F-, *ompT hsdS(rB - mB), dcm+, Tetr gal λ (DE3), endA Hte* [*argU proL Camr*] [*argU ileYleuW Strep/Specr*] | Stratagene, La Jolla, USA |
| ***E. coli* C43 (DE3)** | | (60) |
| **E2407** | pWM321 in *E. coli* JM109 λ pir Amp$^{R}$, Pur$^{R}$ | (57) |
| **E1570** | pET28a(+) in *E. coli* DH5α Kan$^{R}$ | Novagen |
| **K4057** | pRS1214 in *E. coli* DH5α Amp$^{R}$ | This work |
| **K4071** | pRS1223 in *E. coli* DH5α Amp$^{R}$, Kan$^{R}$ | This work |
| **K4076** | pRS1225 in *E. coli* DH5α Kan$^{R}$ | This work |
| **K4078** | pRS1227 in *E. coli* JM109 λ pir Amp$^{R}$, Pur$^{R}$ | This work |
| **K4098** | pRS1240 in *E. coli* DH5α Kan$^{R}$ | This work |
| **K4099** | pRS1240 in *E. coli* BL21 Kan$^{R}$ | This work |
| **Plasmids** | | |
| **pmcl210** | cloning Vector, P15Aori, *lacZ*, *cat* | (61) |
| **pRS207** | pac cassette in pBlueskript SK | (57) |
| **champion pET SUMO** | expression vector, Kan$^{R}$ | Thermo Fischer Scientific, Waltham, USA |
| **pWM321** | shuttle Vector *M. mazei* /*E. coli ori*R6K: pC2A replicon, Amp$^{R}$, Kan$^{R}$ | (62) |
| **pET28a(+)** | expression vector, Kan$^{R}$ | Novagen, Darmstadt, Germany |
| **pET21a(+)** | expression vector, Amp$^{R}$ | Novagen, Darmstadt, Germany |

| | | |
|---|---|---|
| **pCR™II-TOPO®** | Plac, LacZα-ccB, AmpR, KanR, pUC ori | Thermo Fisher Scientific, Waltham, USA |
| **pRS1223** | *pCR™II-TOPO®/MMsORF36 AmpR, KanR* | This work |
| **pRS1225** | *pET28a/MMsORF36 KanR* | This work |
| **pRS1227** | *pWM321/MMsORF36.N* $Pur^R$, $Amp^R$ | This work |
| **pRS1240** | *pETSUMO/MMsORF36.N* $Kan^R$ | This work |
| **pRS1308** | pWM321/MMsORF36 flanking regions/pac casette | This work |
| **pRS1859** | pET21a/*MM733* $Amp^R$ | This work |
