## Supplemental Table 2: Oligonucleotidess for "Small protein mediates inhibition of ammonium transport in Methanosarcina mazei – an ancient mechanism?"

### Table S2: Used Oligonucleotides

| construct | primer | Sequence |
|---|---|---|
| ΔsP36 | DSP35KpnIfor | 5’-GGTACCGCCCTGCTTG |
| | DSP35ecoRIrev | 5’-GTCATCTCCAGTAGAATTCGG |
| | DSP35ecoRIfor | 5’-CTCAGAATTCCTGCTGATCC |
| | DSP35BamHIrev | 5’-CGATTTTCGATGGTGGATCC |
| $His_6$-SUMO-sP36 | sORF36_3_for | 5’-GTGACCATCTGGGAATACGATG |
| | sORF36_3_rev | 5’-AAAAAATTAGAAGGCTGCATCGACATAATC |
| $His_6$-sP36 | sORF36_3_forNdeI | 5’-CATATGGTGACCATCTGGGAATACGATG |
| | sORF36_3_forNdeI | 5’-CATATGAAAAAATTAGAAGGCTGCATCGACATAATC |
| TEV cleavage site | SP36_TEV_rv | 5’-aaataaagattctcgccACCAATCTGTTCTCTGTGAGCCTC |
| | SP36_TEV_fw | 5’-aaataaagattctcgccGGTGTGACCATCTGGGAATACGATG |
