## Supplemental Figures: AlphaFold predictions for "Small protein mediates inhibition of ammonium transport in Methanosarcina mazei – an ancient mechanism?"

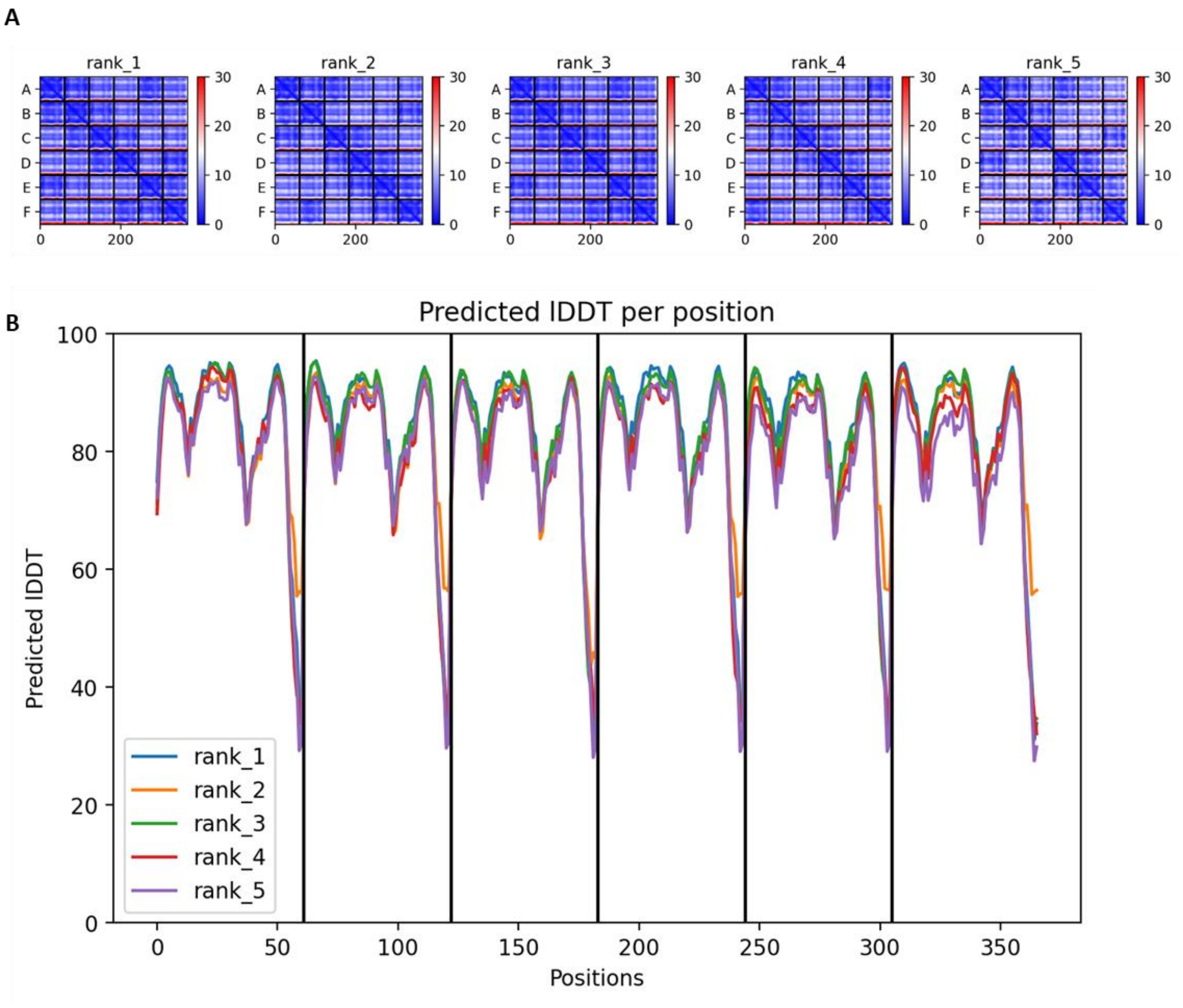


**Supplemental figure S1**. Confidence of the predicted structure of sP36. (A) Predicted aligned error (PAE) and (B) Predicted local distance difference test (IDDT) for the five models generated by AlphaFold2 (50, 51).

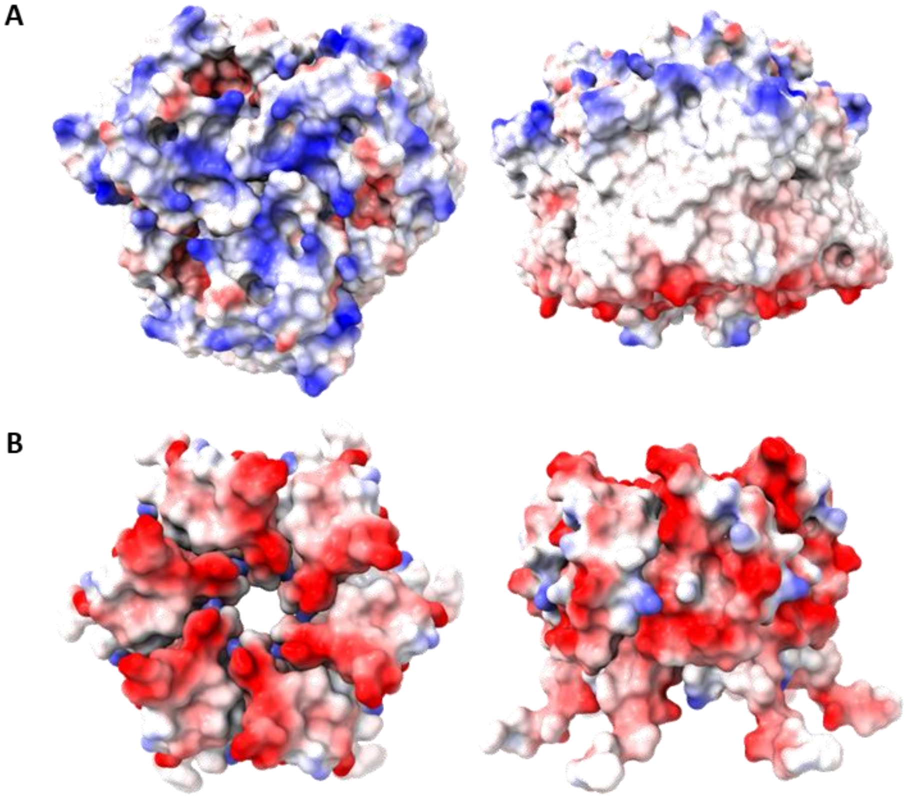


**Supplemental figure S2.** Color-coded representation of the electrostatic potential of the surface of (A) AmtB1 and (B) sP36. The calculation was performed with APBS plug-in implemented in PyMOL (Schrödinger Inc. (2015) The PyMOL Molecular Graphics System. Version 2.0 Schrödinger LLC). Color oscillates from -2.0 (red) to +2.0 (blue) KbT/ec.
