## Supplemental Figure: sP36 deletion southernblot for "Small protein mediates inhibition of ammonium transport in Methanosarcina mazei – an ancient mechanism?"

### Supplemental figures

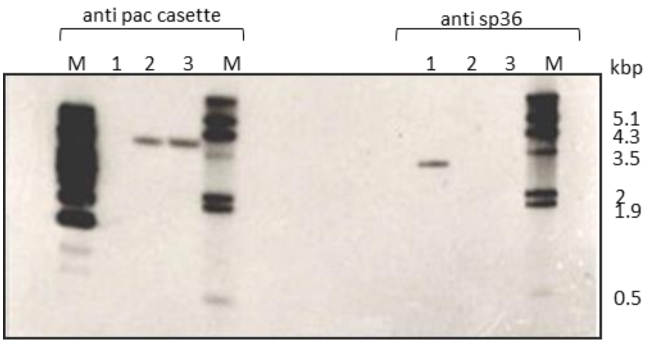


**Supplemental figure S3.** Southern blot of *M. mazei* Δsp36 genomic DNA: to test the success of the sp36 deletion in *M. mazei,* genomic DNA of the new Δsp36 mutant and the wildtype (control) was cleaved by restriction enzyme HindIII and was analyzed with specific probes against sp36 and against the pac casette. **1**: *M. mazei* 3A (wt), **2**: *M. mazei* Δsp36 (clone1), **3**: *M. mazei* Δsp36 (clone2).
